# ATTENTION! The following preprint should no longer be cited as the manuscript and data in their present form are no longer valid!

**DOI:** 10.1101/314898

**Authors:** Nikolaus Brunner, Stephan Groeger, Joao Canelas Raposo, Rupert M. Bruckmaier, Josef J. Gross

## Abstract

Subclinical ketosis (SCK) and periparturient diseases considerably account for economic and welfare losses in dairy cows. The majority of scientific reports investigating the prevalence of SCK and production diseases are based on empirical studies conducted in Western Europe and North America. The present study surveyed the prevalence of SCK and production-related clinical diseases in early lactating cows in various countries across the world other than those in North America and Western Europe. Twelve countries of South and Central America (Argentina, Brazil, Chile, Colombia, Mexico), Africa (South Africa), Asia (Thailand, China), Eastern Europe (Russia, Ukraine), Australia and New Zealand were assessed, and data from a total of 8,902 cows kept at 541 commercial dairy farms were obtained. A minimum of 5 cows per farm were blood sampled and examined once after parturition up to day 21 of lactation. Blood concentration of β-hydroxybutyrate (BHBA) was measured (threshold for SCK: 1.2 mmol/l) and the presence of production-related diseases such as milk fever, retained placenta, mastitis, metritis, displaced abomasum, claw disease and clinical ketosis was recorded. More than 95% of all cows were examined in their second week of lactation. Across all investigated countries, the SCK prevalence was 24.1%, ranging from 8.3% up to 40.1%. The prevalence of production-related diseases detected during the first 21 days of lactation was relatively low (< 5%). Calculated odds ratios did not indicate an elevated risk for production diseases in cows with SCK. Despite differences in production systems across countries and variation between individual farms within a region, the present study data on SCK prevalence align with observations in Western European and North American dairy herds. At the very early stage of sampling and clinical examination for detection of SCK, it cannot be excluded that certain production diseases such as DA, lameness and mastitis have developed later.

## Introduction

At the onset of lactation, dairy cows experience a marked metabolic load due to the prevailing negative energy balance, which makes them susceptible towards infectious and metabolic diseases [1, 2]. Increased concentrations of circulating ketone bodies, predominantly β-hydroxybutyrate (BHBA), without the presence of clinical signs of ketosis are considered as subclinical ketosis (SCK; [3]). The blood BHBA thresholds for SCK diagnosis in literature range between 1.2 and 1.4 mmol/l [4–8]. Although symptoms of clinical ketosis such as reduced milk production, lethargy and loss of appetite are commonly observed at higher concentrations of BHBA (>3.0 mmol/l), some cows, however, may be exposed to high BHBA concentrations without showing any clinical signs, whereas others develop ketosis characteristics already at lower BHBA levels [2, 9, 10].

SCK affects performance and is obviously related to an increased risk of production-related diseases such as clinical ketosis, displaced abomasum (DA), retained placenta, and metritis [11, 12]. Concomitantly, production efficiency decreases (e.g. lower milk production, poor fertility, and increased culling rates) which results in economic losses [11–15]. Serum BHBA concentrations ≥1.2 mmol/l in the first week after calving were associated with a several fold increased risk of subsequently developing DA and metritis, respectively [11]. Ospina et al. [16] found increased risk ratios of 4.9 for clinical ketosis, 2.3 for metritis and 6.9 for DA when plasma BHBA concentrations exceeded a threshold of 1.0 mmol/l during days 3 to 14 post partum. A negative impact of SCK on milk yield of early lactating cows was reported by several authors [10–12, 17]. Furthermore, cows with BHBA concentrations above 1.0 mmol/l in the first week after calving were significantly less likely to be diagnosed pregnant after first insemination [13]. The economic impact of SCK is therefore indisputable, although exact figures quantifying the actual costs of SCK are difficult to collect. Based on an earlier investigation on Canadian dairy farms including more than 2,600 cows, McLaren et al. [18] estimated that a reduction of 1% in SCK incidence would amount to a saving of 584 USD per year. Recently, McArt et al. [14] calculated the average total costs of hyperketonemia (blood BHBA concentrations ≥1.2 mmol/l) to be 289 USD per case diagnosed.

Prevalence of SCK in the existing literature ranges between 10 and 40% in early lactation with highest values occurring within the first three weeks of lactation [11, 17, 19]. Most of these studies, however, included only a small number of dairy herds and cows (mainly research stations), and were performed in North America and Western Europe [5–8, 12].

The objective of the present study was to survey the prevalence of SCK and the concomitant occurrence of health disorders in early lactating dairy cows in various countries across the world other than those in North America and Western Europe. Dairy farms in 12 countries of South and Central America (Argentina, Brazil, Chile, Colombia and Mexico), Africa (South Africa), Asia (Thailand, China), Eastern Europe (Russia, Ukraine), Australia and New Zealand were investigated. To our knowledge, this is the first large-scale approach to investigate the prevalence of SCK beyond Western Europe and North America.

## Materials and methods

### Ethics statement

All procedures in terms of blood sampling and veterinary examinations followed the animal care and welfare legislation criteria of the involved countries. The approval was provided by the local responsible institutions of the following countries: Argentina, Australia, Brazil, Chile, China, Colombia, Mexico, New Zealand, Russia, South Africa, Thailand and Ukraine.

### Animals, blood sampling and analysis

The study was conducted from June 2011 to September 2013 at different commercial dairy farms in 12 different countries worldwide (Argentina, Australia, Brazil, Chile, China, Colombia, Mexico, New Zealand, Russia, South Africa, Thailand and Ukraine). Breeds included were dairy breeds (e.g. predominantly Holstein Friesian, Jersey), local crossbred lines (e.g. Girolondo in Brazil, Simmental x Red Steppe in Russia) and others (e.g. Simmental). Management systems varied from grazing systems to indoor housing of animals throughout the year. In total, 8,902 early lactating dairy cows housed on 541 farms were evaluated. At least five randomly selected cows of a herd in early lactation (day 2 to 21 post partum (p.p.)) inconspicuous of obvious health disorders and without any previous treatments were clinically examined and blood was sampled for diagnostic purposes and health monitoring during regular herd health management visits by veterinarians in the field. Each cow was only tested once and no further restrictions in selection criteria such as parity number, breed, or performance were implied. In all cows, blood samples were taken from the coccygeal vein, and BHBA concentrations were directly determined on site using a handheld meter (Precision Xceed, Abbott Diabetes Care Inc., Alameda, CA, USA), which was previously validated for the use in cows [20, 21]. Concomitantly, each sampled cow was clinically examined for the presence of metabolic and infectious health disorders (milk fever, retained placenta, displaced abomasum (DA), claw disease, mastitis, metritis and clinical ketosis (CK)) following standardized diagnostic definitions (Table 1).

**Table 1.**
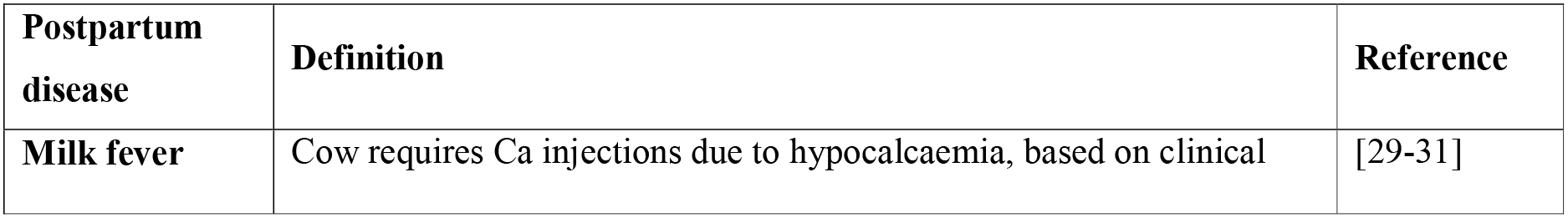
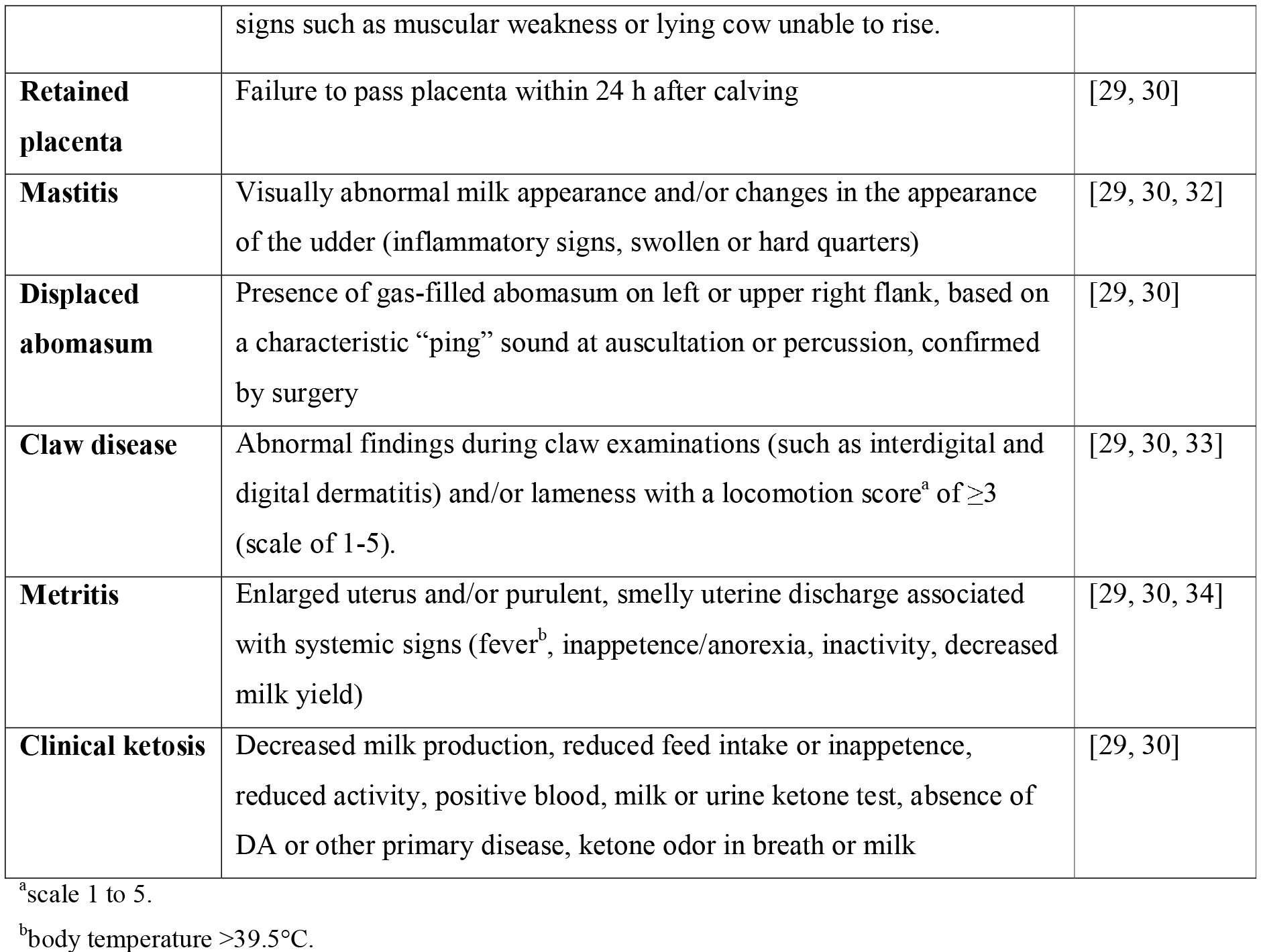
Definitions used for disease diagnosis during veterinary examinations of dairy cows in early lactation.

### Statistical analysis

Cows sampled between day 2 and 21 p.p. were categorized “SCK negative” if blood BHBA concentrations were <1.2 mmol/l, or “SCK positive” when BHBA concentrations were ≥1.2 mmol/l). The prevalence of SCK was calculated as proportion of animals with BHBA concentrations ≥1.2 mmol/l relative to all investigated cows. Prevalence data were estimated at country and average global level. In addition, overall and country-specific prevalence of postpartum diseases was calculated. The occurrence of postpartum diseases in relation to SCK was evaluated by odds ratio (OR) analyses using the statistical package TESTIMATE (version 6.5, idv-Data Analysis & Study Planning, Krailling, Germany) according to following formula: OR = (p_1_/(1-p_1_)) / (p_2_/(1-p_2_)), where p_1_ indicates the probability of postpartum diseases under SCK conditions and p_2_ the probability of postpartum diseases under non-SCK conditions.

## Results

Data analyses were based on a total of 8,902 dairy cows from 541 different commercial dairy farms. The number of farms investigated per country and their herd size ranged from two farms with 83 cows in Thailand to 102 farms housing 2,989 cows in New Zealand. On average, cows of the present dataset were in their 3^rd^ lactation (range of parity number: 1 - 14, 26.9% primiparous and 73.1% multiparous dairy cows; Table 2). The majority (95.1%) of the cows was examined and sampled between days 7 and 15 p.p..

**Table 2.**
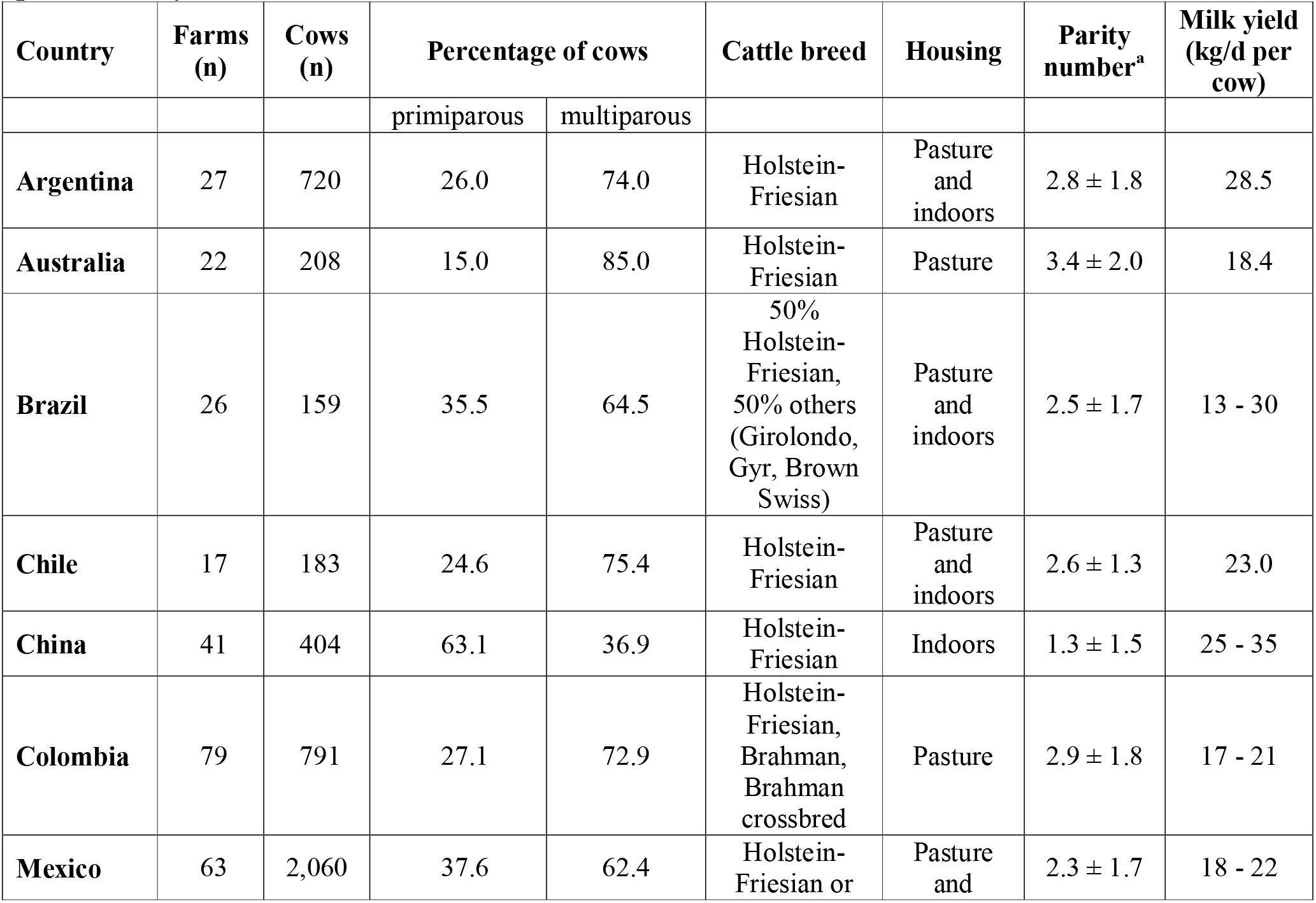
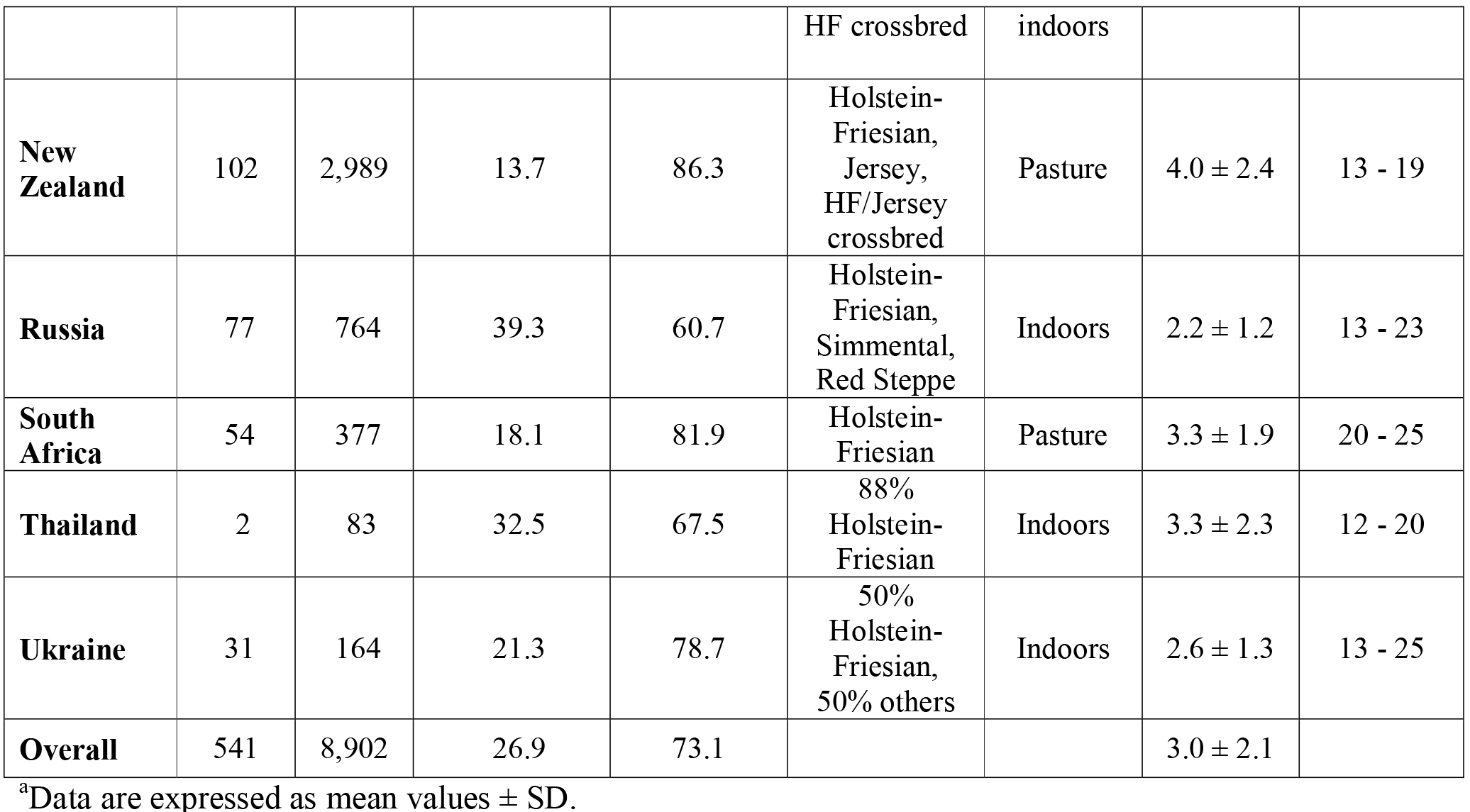
Information on animals and farms of the different participating countries in the present study.

| Country | Farms (n) | Cows (n) | Percentage of cows |  | Cattle breed | Housing | Parity number<sup>a</sup> | Milk yield (kg/d per cow) |
| --- | --- | --- | --- | --- | --- | --- | --- | --- |
|  |  |  | primiparous | multiparous |  |  |  |  |
| Argentina | 27 | 720 | 26.0 | 74.0 | Holstein-Friesian | Pasture and indoors | 2.8 ± 1.8 | 28.5 |
| Australia | 22 | 208 | 15.0 | 85.0 | Holstein-Friesian | Pasture | 3.4 ± 2.0 | 18.4 |
| Brazil | 26 | 159 | 35.5 | 64.5 | 50% Holstein-Friesian, 50% others (Girolondo, Gyr, Brown Swiss) | Pasture and indoors | 2.5 ± 1.7 | 13 - 30 |
| Chile | 17 | 183 | 24.6 | 75.4 | Holstein-Friesian | Pasture and indoors | 2.6 ± 1.3 | 23.0 |
| China | 41 | 404 | 63.1 | 36.9 | Holstein-Friesian | Indoors | 1.3 ± 1.5 | 25 - 35 |
| Colombia | 79 | 791 | 27.1 | 72.9 | Holstein-Friesian, Brahman, Brahman crossbred | Pasture | 2.9 ± 1.8 | 17 - 21 |
| Mexico | 63 | 2,060 | 37.6 | 62.4 | Holstein-Friesian or | Pasture and | 2.3 ± 1.7 | 18 - 22 |
|  |  |  |  |  | HF crossbred | indoors |  |  |
| **New Zealand** | 102 | 2,989 | 13.7 | 86.3 | Holstein-Friesian, Jersey, HF/Jersey crossbred | Pasture | 4.0 ± 2.4 | 13 - 19 |
| **Russia** | 77 | 764 | 39.3 | 60.7 | Holstein-Friesian, Simmental, Red Steppe | Indoors | 2.2 ± 1.2 | 13 - 23 |
| **South Africa** | 54 | 377 | 18.1 | 81.9 | Holstein-Friesian | Pasture | 3.3 ± 1.9 | 20 - 25 |
| **Thailand** | 2 | 83 | 32.5 | 67.5 | 88% Holstein-Friesian | Indoors | 3.3 ± 2.3 | 12 - 20 |
| **Ukraine** | 31 | 164 | 21.3 | 78.7 | 50% Holstein-Friesian, 50% others | Indoors | 2.6 ± 1.3 | 13 - 25 |
| **Overall** | 541 | 8,902 | 26.9 | 73.1 |  |  | 3.0 ± 2.1 |  |

<sup>a</sup>Data are expressed as mean values ± SD.

### Concentration of blood BHBA and subclinical ketosis

Blood BHBA concentrations observed between day 2 and day 21 p.p. were 1.0 ± 0.8 mmol/l, ranging from an average value of 0.7 mmol/l (Australia, Brazil, Colombia and Russia) up to 1.5 mmol/l (Ukraine). In two thirds of the investigated countries, mean BHBA concentrations were below 1.0 mmol/l (0.7 - 0.9 mmol/l) while in one third of the countries (namely China, New Zealand, Thailand and Ukraine) mean BHBA concentrations of 1.0 mmol/l and above were observed (Table 3). Blood BHBA concentrations of 3.0 mmol/l and beyond during the first 21 days of lactation were predominately observed in China and Ukraine (Table 3). Overall, SCK (BHBA concentrations ≥1.2 mmol/l) was diagnosed in 24.1% of all cows examined, ranging from 8.3% (Colombia) up to 40.1% (New Zealand; Fig 1). In only two (Australia and Colombia) out of the 12 investigated countries, SCK prevalence on the participating farms was less than 10%. In four countries (Brazil, Chile, Mexico, Russia) SCK prevalence varied between 10.7% and 14.8%, whereas in the remaining countries (Argentina, China, New Zealand, South Africa, Thailand, Ukraine) SCK prevalence was above 15% (between 17.0% and 40.1%). More than 80% of all cows diagnosed SCK were multiparous. In contrast, we observed SCK occurring more frequently in primiparous cows in three countries (Chile, South Africa, Ukraine; Fig 2).

**Table 3.** Blood BHBA concentrations in early lactating dairy cows (2- 21 days postpartum) in different countries and proportion of cows with highly elevated BHBA concentrations.

| Country | BHBA concentration (mmol/l) | | | Cows with BHBA $\geq 3.0$ mmol/l | |
| --- | --- | --- | --- | --- | --- |
| | n | mean ± SD | range | n | % |
| Argentina | 720 | 0.9 ± 0.9 | 0.1 - 6.2 | 36 | 5.0 |
| Australia | 208 | 0.7 ± 0.7 | 0.1 - 7.0 | 4 | 1.9 |
| Brazil | 159 | 0.7 ± 0.6 | 0.1 - 4.8 | 3 | 1.9 |
| Chile | 183 | 0.8 ± 0.7 | 0.0 - 4.6 | 3 | 1.6 |
| China | 404 | 1.2 ± 1.0 | 0.0 - 5.8 | 33 | 8.2 |
| Colombia | 791 | 0.7 ± 0.4 | 0.1 - 4.2 | 4 | 0.5 |
| Mexico | 2,060 | 0.8 ± 0.7 | 0.0 - 6.0 | 53 | 2.6 |
| New Zealand | 2,990 | 1.2 ± 0.7 | 0.0 - 8.0 | 101 | 3.4 |
| Russia | 764 | 0.7 ± 0.7 | 0.1 - 6.4 | 18 | 2.4 |
| South Africa | 376 | 0.9 ± 1.0 | 0.1 - 7.2 | 20 | 5.3 |
| Thailand | 83 | 1.0 ± 0.8 | 0.1 - 5.8 | 2 | 2.4 |
| Ukraine | 164 | 1.5 ± 1.5 | 0.0 - 6.7 | 22 | 13.4 |
| Overall | 8,902 | 1.0 ± 0.8 | 0.0 - 8.0 | 299 | 3.4 |

**Fig 1. Prevalence of subclinical ketosis worldwide.** Prevalence of subclinical ketosis (SCK, blood BHBA concentrations ≥1.2 mmol/l) in early lactating dairy cows studied in 12 countries worldwide.

**Fig 2. Subclinical ketosis in primi- and multiparous dairy cows.** Occurrence of subclinical ketosis (SCK, blood BHBA concentrations ≥1.2 mmol/l) in primiparous and multiparous cows during early lactation in different countries worldwide.

### Prevalence of concomitant production diseases and their association with SCK

The overall prevalence of production diseases was 4.3% for milk fever, 4.0% for retained placenta, 3.4% for mastitis, 1.7% for claw disease and 5.3% for metritis (Table 4). Displaced abomasum (DA) and clinical ketosis showed a very low prevalence (0.3% and 7%, respectively). During the first 21 days p.p., DA was not seen in 50% (6/12) of the investigated countries and CK was diagnosed in only 7 out of 12 countries. We further investigated the relationship of SCK with the concomitant presence of health disorders in early lactation by performing an odds ratio analysis. However, due to the generally low prevalence of diseases, odds ratios (OR) were <1, and an increased risk of disease in association with SCK could not be identified in the present study (data not shown). The highest odds ratio was calculated for clinical ketosis as cows diagnosed SCK had a 1.062 higher probability of having subsequently clinical ketosis.

**Table 4.** Prevalence of production diseases between 2 and 21 days postpartum.

| Country | Milk fever |  | Retained placenta |  | Mastitis |  | DA<sup>a</sup> |  | Claw disease |  | Metritis |  | CK<sup>b</sup> |  |
| --- | --- | --- | --- | --- | --- | --- | --- | --- | --- | --- | --- | --- | --- | --- |
|  | % | n | % | n | % | n | % | n | % | n | % | n | % | n |
| Argentina | 0.7 | (5/720) | 2.1 | (15/720) | 5.1 | (37/720) | 1.0 | (7/720) | 2.2 | (16/720) | 15.7 | (113/720) | 4.0 | (29/720) |
| Australia | 10.1 | (21/208) | 7.2 | (15/208) | 3.8 | (8/208) | 0 | (0/208) | 0.5 | (1/207) | 6.3 | (13/208) | 1.9 | (4/208) |
| Brazil | 0 | (0/159) | 8.8 | (14/159) | 3.8 | (6/159) | 0.6 | (1/159) | 1.3 | (2/159) | 0 | (0/159) | 0 | (0/159) |
| Chile | 7.1 | (13/182) | 5.5 | (10/183) | 9.3 | (17/183) | 1.1 | (2/183) | 5.5 | (10/183) | 9.8 | (18/183) | 2.2 | (4/183) |
| China | 1.7 | (7/403) | 4.0 | (16/404) | 2.0 | (8/404) | 2.7 | (14/404) | 1.0 | (4/404) | 3.7 | (15/404) | 1.2 | (5/404) |
| Colombia | 1.5 | (12/791) | 4.3 | (34/791) | 4.6 | (36/791) | 0.1 | (1/791) | 1.0 | (8/791) | 0.9 | (7/791) | 0 | (0/791) |
| Mexico | 0.6 | (13/2060) | 4.7 | (97/2060) | 1.4 | (29/2060) | 0.2 | (4/2060) | 1.3 | (27/2060) | 5.7 | (118/2059) | 0.6 | (12/2060) |
| New Zealand | 0.7 | (20/2990) | 0.2 | (7/2990) | 1.2 | (36/2990) | 0 | (0/2990) | 0.1 | (3/2990) | 0.3 | (10/2990) | 0.1 | (2/2989) |
| Russia | 37.4 | (286/764) | 17.0 | (130/764) | 14.9 | (114/764) | 0 | (0/764) | 10.5 | (80/764) | 13.5 | (103/764) | 0.9 | (7/764) |
| South Africa | 0.8 | (3/377) | 1.3 | (5/377) | 0.8 | (3/377) | 0 | (0/377) | 0 | (0/377) | 6.6 | (25/377) | 0 | (0/377) |
| Thailand | 1.2 | (1/83) | 0 | (0/83) | 4.8 | (4/83) | 0 | (0/83) | 1.2 | (1/83) | 19.3 | (16/83) | 0 | (0/83) |
| Ukraine | 0.6 | (1/164) | 7.3 | (12/164) | 3.7 | (6/164) | 0 | (0/164) | 0.6 | (1/164) | 19.5 | (32/164) | 0 | (0/164) |
| Overall | 4.3 | (382/8901) | 4.0 | (355/8903) | 3.4 | (304/8903) | 0.3 | (26/8903) | 1.7 | (153/8902) | 5.3 | (470/8902) | 0.7 | (63/8902) |
| Range | 0 - 37.4 % |  | 0 - 17.0 % |  | 0.8 - 14.9 % |  | 0 - 2.7 % |  | 0 - 10.5 % |  | 0 - 19.5 % |  | 0 - 4.0 % |  |

<sup>a</sup>DA = displaced abomasum.
<sup>b</sup>CK = clinical ketosis.

## Discussion

In the last three decades, research results addressing SCK and periparturient diseases represented primarily published data from the United States, Canada and Western Europe [6, 12, 22]. The majority of reports usually based on investigations conducted in regional and small-sized dairy farms of individual countries [8, 16, 18, 19]. Studies involving a higher number of animals and covering wider geographic areas were presented earlier by Chapinal et al. [23] and Suthar et al. [6], but still considered only SCK occurrence within Europe and the United States, respectively. Different study designs and scopes (such as pre- or postpartum reclassification of animals, herd or cow level investigations, sampling of blood, milk or urine, and defining SCK at various BHBA thresholds) do not always allow a straight forward comparison of literature data across farms, regions and countries, as recently described in a meta-analysis evaluating 23 SCK publications by Raboisson et al. [22]. Furthermore, data on global SCK prevalence obtained by a standardized protocol for sampling and defining SCK are scarce. Therefore, we have investigated the prevalence of SCK and typical production-related disorders in early lactating dairy cows in various countries across the world other than those in North America and Western Europe. By using the same validated tool for on-site BHBA testing in all cows, diagnosing SCK at the generally accepted blood BHBA cut-off concentration of 1.2 mmol/l and sampling cows in the very early lactation (days 2 to 21 p.p.) make the results presented here comparable to studies performed in other countries.

The overall SCK prevalence of the present survey was in a comparable range to observations from a European evaluation of Suthar et al. [6]. In addition, the variation in the global SCK rates presented here highly scatters (approximately 8 - 40%) and is similar to findings reported for US and Europe [6, 8, 12]. The majority of our investigated cows with SCK were multiparous confirming the assumptions that the risk of developing SCK increases with parity number as milk production rises concomitantly [24]. Based on an analysis of a variety of publications, Oetzel [19] considered a prevalence rate of 15% representative for SCK in dairy herds, but emphasized to consider a SCK prevalence of 10% already as an alert level. In the present study only 2 out of 12 countries (Australia and Colombia) had an average SCK prevalence of less than 10%, whereas the remaining countries had higher rates. Four countries (China, New Zealand, Ukraine, and Thailand) revealed even SCK rates up to 40%. However, this does not exclude that SCK may also occur at higher rates in the countries with low SCK prevalence in the present study which might be attributed to the random selection of farms and animals. In general SCK in early lactating dairy cows is globally present. The highest average SCK prevalence of all countries was observed in New Zealand. One explanation might be that dairy cows in New Zealand are mostly kept under extensive pasture based production conditions with low or even zero concentrate supplementation [7]. Along with the genetic progress in dairy cow breeding, milk production in export-oriented countries such as New Zealand likely increased and aggravated the energy deficiency in early lactating cows [2, 25]. In another study, Compton et al. [7] observed a lower SCK prevalence of approximately 17% in New Zealand dairy cows, which would be still in the upper range when compared to the present evaluations. The difference just implies the possible variation due to the selection of farms and animals.

Interestingly, the occurrence of postpartum diseases of the present study was much lower compared to literature reports. Our observations on average prevalence rates of retained placenta, mastitis, claw diseases and metritis were approximately half of the values reported in a European study by Suthar et al. [6]. At first sight, these results were surprising considering the fact that SCK prevalence rates of this investigation were in agreement with other reports published. The cows diagnosed SCK in the present study did obviously not develop health disorders at this early sampling stage. However, results of the concomitant presence of SCK and further production diseases are inconsistent. While we observed the highest SCK prevalence on farms in New Zealand, postpartum diseases occurred only at less than 1% in cows diagnosed SCK. On the other hand, the SCK prevalence data in Russia were only moderately high (14.1%), but many health disorders were diagnosed at the same time (milk fever 37.4%, retained placenta 17.0%, mastitis 14.9%, claw disease 10.5%). In a recent study by Zbinden et al. [10], high-yielding cows fed only herbage showed blood BHBA concentrations approaching that representative for clinical ketosis, however, not necessarily followed by the occurrence of diseases. Irrespective of the concomitant presence of production diseases in cows with SCK, elevated BHBA concentrations ≥1.2 mmol/l undoubtedly increase the risk of subsequently developing further health disorders [11, 16] at simultaneously reduced performance in terms of milk yield and reproduction [11–13, 17]. Management and environmental conditions are crucial factors that affect the ability of cows to cope successfully with the imposed metabolic load. These circumstances cannot be effectively recorded in a field study like the present one. Furthermore, one might speculate on regional differences in terms of breed and feeding system affecting the development of health disorders. The occurrence of DA was reported to be relatively rare in dairy cattle in New Zealand and Australia [7, 26], which is assumed to be associated with the predominantly extensive farming on pasture where important risk factors for DA such as low locomotion activity are not important. However, prevalence of DA concomitant with SCK in the present study was in particular low in countries with a more intensive milk production in confined housing systems (e.g. China, Russia, Thailand, and Ukraine). Similar observations were made for mastitis and the occurrence of lameness. Although lameness may also occur in cattle on pasture, it appears to be less frequent and to be usually based on a different aetiology [27, 28]. In contrast, we observed that mastitis prevalence in cows with SCK can be elevated independent if cows were kept on pasture or indoors.

## Conclusions

In conclusion, findings of the present study showed that SCK in dairy cows is a global issue and can be observed at various rates in many countries over the world. Possibly due to the diagnosis of SCK and clinical examination at a very early stage of lactation, concomitant postpartum diseases in cows diagnosed for SCK have not yet developed at this stage. An appropriate management is crucial for avoiding a deterioration of animals’ metabolic status with the associated negative impacts on health and performance. The results of the present study support the hypothesis that the association between SCK and postpartum diseases is complex and that based on an occasional single determination of blood BHBA concentrations alone, a reliable and robust prediction of further production diseases is not possible.

## Acknowledgements

Authors are grateful to all involved veterinarians, farmers and staff on the different farms and countries. Furthermore we would like to thank Mrs. Marion Ocak for her contribution to the statistical analysis and Mrs. Tanja Knoppe for data processing and literature research. This study was supported by Bayer Animal Health GmbH, 51368 Leverkusen, Germany.

